## Supplemental Information for "A structural dynamics model for how CPEB3 binding to SUMO2 can regulate translational control in dendritic spines"

### Definition of the structural similarity metric Q value

As a metric of structural similarity for 2 configurations, the Q value ranges from 0 to 1, lower values reflecting low similarity while Q=1 indicates the structures are nearly exactly the same. The definition of the mutual Q for two structure A, B is shown below,

$$Q = \frac{1}{N} \sum_{ij} e^{-\frac{(r_{ij}^{A} - r_{ij}^{B})^2}{2\delta_{ij}^{2}}}$$

where N is the total number of residue pairs considered in Q calculation, i, j are residue indices and $r_{ij}^{A}$ and $r_{ij}^{B}$ are the distances between residue i and residue j in structures A, B respectively. For intra-molecular residue pairs, $\delta_{ij} = |i - j|^{0.15}$, while for inter-molecular residue pairs, $\delta_{ij}^{2}$ is set to a constant value, 5 $A^2$. We included all the inter-molecular and inter-domain residue pairs for calculating mutual Q in clustering analysis and $Q_f$ in free energy analysis. For the Q values of RRM1-SIM/SUMO2 when comparing with the canonical SIM/SUMO2 template, we only considered inter-molecular residue pairs which form contacts in the template structure or in the predicted structure. In this work, we chose a distance threshold of 12 A to define contacts between residues.

### Introduction to the AWSEM-3SPN2 force field

In this study, all simulations were carried out using the AWSEM(-3SPN2) force field. The AWSEM force field is a predictive coarse-grained model for proteins, whose Hamiltonian $V_{AWSEM}$ is shown below,

$$V_{AWSEM} = V_{backbone} + V_{contact} + V_{burial} + V_{DH} + V_{HB} + V_{FM},$$

where $V_{backbone}$ is a peptide-like backbone term; $V_{contact}$ is a tertiary residual interaction term; $V_{burial}$ is a many body burial term; $V_{DH}$ is the Debye Huckel electrostatic term; $V_{HB}$ is a beta-sheet favored, hydrogen bond term; $V_{FM}$ is the associative fragment memory term, a bioinformatical term guiding the local-in-sequence interactions. In all simulations, we used the PDB structure 2N1W as the only memory for SUMO2 protein. We used the PDB structure 2MKJ to model the structure of free state CPEB3-RRMs and used the PDB structure 2M13 to model the partial structure of CPEB3-ZnF via Modeller. These two structures are then used as the only memories for RRMs and partial ZnF. The fragment memories for the rest of the ZnF were obtained from the known protein data bank using a sequence alignment algorithm.

For simulations containing the RNA fragment, we used the 3SPN2 force field which is a coarse grained model originally developed for simulating DNA, as the model for the RNA by regarding Uracil (U) as the base particle thymine (T) in 3SPN2:

$$V_{3SPN2} = V_{backbone} + V_{stacking} + V_{BP} + V_{CS} + V_{excl} + V_{DH},$$

where $V_{backbone}$ is a bonded term connecting 3-beads nucleotides backbone; $V_{stacking}$ is a stacking term between consecutive nucleotides; $V_{BP}$ is a base pairing term between complementary nucleobases; $V_{CS}$ is a cross stacking term between neighboring base pairs; $V_{excl}$ is an exclusion term; $V_{DH}$ is the Debye Huckel electrostatic term. The interaction between the protein and the nucleotides is modelled by an exclusion term and the Debye Huckel electrostatic term in AWSEM-3SPN2 force field. A more detailed introduction to the AWSEM-3SPN2 force field can be found in previous papers[1, 2, 3, 4]. A debye length of 10 A is used for all Debye Huckel electrostatic terms. This value corresponds to a typical physiological solution condition when temperature T=300K and ionic strength equals 0.1 M, using a dielectric constant for water $\epsilon = 80$[5]. Original AWSEM-3SPN2 only assigns charges to the phosphate particle (-0.6) and four types of residues (Arg:+1, Lys:+1, Asp:-1, and Glu:-1). Here additional charges were applied to the two zinc coordination sites include Zn1 (Cys654, Cys657, Cys681, Cys684) and Zn2 (Cys671, Cys676, His689, His697) to model the two zinc ions (+0.5 for each coordination residue) . The first nucleotide of the nucleic acid chains in 3SPN2 model lacks the phosphate particle, which is a proper approximation for very long nucleic acid chains. However, our simulations used a short RNA chain with a length of five nucleotides. Therefore, we assigned an additional charge of -0.6 to the sugar particle of the first nucleotide of our RNA chains. For the electrostatic interactions between protein and RNA, we modified the effective charge of each nucleotide into -1.0. Other parameters are set to the default values in AWSEM-3SPN2 (openmm version) and can be found at https://github.com/npschafer/openawsem and https://github.com/cabb99/open3spn2.

### Binding energy for SIM/SUMO2 complex

The binding energy $E_{binding}$ for the SIM/SUMO2 complex is calculated using the equation:

$$E_{binding} = E_{SIM/SUMO2} - E_{SIM} - E_{SUMO2},$$

where $E_{SIM/SUMO2}$, $E_{SIM}$, $E_{SUMO2}$ are the AWSEM energies for the SIM/SUMO complex, the SIM peptide and the SUMO2 protein separately. Here we used a complex structure (PDB ID: 6JXW) of a canonical SIM peptide (12 amino acids) and SUMO2 protein as the template and changed the sequence of the SIM peptide into the truncated CPEB3 sequence. The truncated CPEB3 sequence (12-amino acids) scanned over full length CPEB3 sequence with a 11-amino acids overlap between neighboring peptide sequences to calculate the binding energy of CPEB3 peptides with SUMO2 (Fig. S2).

### Additional potentials

To model the two zinc ions coordinating with ZnF domain, we have assigned positive charges to coordination residues. However, this modification of charges can not take into account all the structural effects of coordinated zinc ions. To confine the structure of ZnF, we applied an AMHGo potential ($V_{AMHGo}$) to all simulations containing the ZnF domain:

$$V_{AMHGo} = -k_{AMHGo} \sum_{ij} e^{-\frac{(r_{ij}^A - r_{ij}^B)^2}{2\delta_{ij}^2}},$$

where $k_{AMHGo} = 0.3 kcal/mol$ and other parameters are the same as corresponding parameters in Q value definition. Here structure A represents the ZnF structure during simulations and structure B is the Modeller structure of ZnF built from the PDB structure 2M13. We only considered the residue pairs with a distance closer than 8 A in the template structure.

To guide the formation of the inter-molecular beta sheet between RRM1-SIM and SUMO2-$\beta$2, we applied a spring potential ($V_{rbias}$) between these two groups of residues for a biased simulation:

$$V_{rbias} = \frac{1}{2} k_{rbias} (r - r_0)^2,$$

where $k_{rbias} = 0.1 kcal/mol/A^2$, $r_0 = 15A$ and r is the distance between the mass centers of RRM1-SIM (P484-W489) and SUMO2-$\beta$2 (G27-R36).

To run the umbrella sampling simulations for studying the closure motion of RRMs, we used a biasing potential for the reaction coordinate $Q_f$:

$$V_{qbias} = \frac{1}{2} k_{qbias} (Q_f - Q_0)^2,$$

where $k_{qbias} = 10000 kcal/mol$, $Q_0$ is the minimal for a simulation window and $Q_f$ is the inter-domain Q value for current structure corresponding to the NMR structure of free RRMs.

To simulate the specific binding between CPEB3-RRMs and its target mRNA, we applied a sequence-specific contact term $V_{ssc}$ between the protein and the RNA for all simulations containing RNA:

$$V_{ssc} = -k_{ssc} \sum_{ij} \gamma_{ij} e^{-\frac{(r_{ij} - r_{ij}^B)^2}{2\delta^2}},$$

where $k_{ssc} = 0.8 kcal/mol$, $\delta = 1.5A$, $r_{ij}, r_{ij}^B$ are the distance between residues i and nucleotide j in current structure and the NMR structure of RNA-bound RRMs. $\gamma_{ij}$ is the weight for each residue-nucleotide pair and equals $ln(C_{ij})$, where $C_{ij}$ is the atomic contact number between residues i and nucleotide j in the NMR structure of RNA-bound RRMs. Here we used a distance threshold of 4.5 A to define atomic contacts and only considered residue-nucleotide pairs having no less than 14 atomic contacts in NMR structure ($C_{ij} \geq 14$). Given these parameter values, the RNA-binding affinity of RRMs calculated from our free energy profile ($\sim$ 8 kacl/mol) is comparable to the experimental value (6.4 $\sim$ 6.9 kcal/mol).

To calculate the free energy profiles of RNA dissociation for protein/RNA complex systems, we ran a two-dimentional umbrella sampling for each system. The major reaction coordinate is the value of $V_{ssc}$, corresponding to the biasing potential:

$$V_{sscbias} = \frac{1}{2} k_{sscbias} (V_{ssc} - V_0)^2,$$

where $k_{sscbias} = 1 mol/kcal$ and $V_0$ is the minimal for a simulation window. For simulation windows with $V_0$ values close to 0, an additional biasing potential $V_{Rbias}$ was applied to enhance sampling along the order parameter R, the distance between RNA and RNA-binding pocket in RRMs:

$$V_{Rbias} = \frac{1}{2} k_{Rbias} (R - R_0)^2,$$

where $k_{Rbias} = 0.1 kcal/mol/A^2$, $R_0$ the minimal for a simulation window and R is the distance between the mass centers of RNA chain and RNA-binding pocket (F462, G465, P490, Y504, F506, K541, Q544, F570, K609, K645).

### Simulation methods

All simulations were run in the openMM platform with the Langevin integrator at a constant temperature 300 K. We used 2 fs as the simulation time step and 1 ps as the damping time. For all umbrella sampling simulations, we output one sampling structure every 5000 steps.

#### Equilibrum simulation of free RRMs

We ran 20 independent equilibrium simulations of free RRMs, starting from the Modeller structure built from PDB structure 2MKJ. Each simulation is run for $6 \times 10^6$ steps and outputs one sampling structure every 10000 steps. All sampled structures in 20 trajectories were then used for the cartesian principal component analysis (PCA). We first superposed the beta sheet of RRM1 domain of each sampled structure with that of the initial structure, using several residues of the RRM1 beta sheet (V463-P468,E525,D526,Y530,Q544-P547) as targets. Then updated 3-dimentional coordinates of all the residues are recorded as PCA input data. The first principal component PC0 is then used as the order parameter to describe RRMs closure motion. The final frames of the 20 simulations are considered as predicted structures and were clustered into 3 subgroups (Fig. S3).

### Equilibrum simulation of full length RBD

By fusing the ZnF domain to the C-terminal of the three representative RRMs structures shown in Fig. S3, we obtained three initial structures for full length RBD simulation. We ran 20 independent equilibrium simulations for each initial structure (60 simulations overall). Each simulation was run for $6 \times 10^6$ steps and the final frame is considered as the single predicted structure for full length RBD.

### Structural prediction to the SUMO2/RBD complex

Using the NMR structure of SUMO1/CBP-ZnF complex (PDB ID: 2N1A) as a template, we can dock the SUMO2 protein to the ZnF of the 60 predicted structures of full length RBD and obtain 60 initial structures for the SUMO2/RBD complex structure prediction. For each initial structure, a biased AWSEM simulations with an additional potential $V_{rbias}$ was run for $6 \times 10^6$ steps. After turning the $V_{rbias}$ term off, the last frame of the biased simulation was relaxed for $2 \times 10^7$ steps. The final structures of the relaxed simulations were considered as candidates for further evaluation and selection.

### Free energy profiles for RNA dissociation

For the RRMs/RNA system, we used the Modeller structure built from the NMR structure of RNA-bound RRMs (PDB ID: 2MKI) as the initial structure. The RNA sequence used in all simulations is also the same as the sequence in PDB structure 2MKI: 5'-CUUUA-3'. For the RBD/RNA system, we attached the ZnF domain to the C-terminal of RRMs in the RRMs/RNA Modeller structure to obtain the initial structure. For the SUMO2/RBD/RNA system, we first docked the RNA chain to the RNA-binding pocket in the predicted SUMO2/RBD complex. Then we ran a biased equilibrium simulation for $8 \times 10^6$ steps with the $V_{sccbias}$ term on. In this simulation, we set $k_{sscbias} = 5mol/kcal$ and $V_0 = -35kcal/mol$ so that the sequence specific protein/RNA contacts were correctly formed in the final structure, which was then used as the initial structure for later umbrella sampling simulations.

For all these RNA-bound systems, we applied a 2D umbrella sampling method to archieve sufficient sampling using both $V_{ssc}$ and $R$ as two biasing reaction coordinates. The biasing potentials $V_{sscbias}$ and $V_{Rbias}$ are mentioned above. We first turned off the $V_{Rbias}$ term and set 33 simulation windows with $V_0$ evenly distributing from -35 kcal/mol to -3 kcal/mol. We also turned on the $V_{Rbias}$ term and set 9 simulation windows with $R_0$ evenly distributing from 1 nm to 5 nm when $V_0$ equals -2, -1 or 0 kcal/mol. (So there are 60 simulation windows overall.) Each simulation lasts $8 \times 10^6$ steps. The first $1 \times 10^6$

steps are treated as equilibration and are then discarded in later analysis. The free energy profiles are calculated by 2D-WHAM analysis[6] and projected into another order parameter, PC0.

#### Free energy profiles for the closure motion of RRMs

For all the systems containing the RBD (including RBD, SUMO2/RBD shown in Fig. 4 and the RBD/RNA, SUMO2/RBD/RNA shown in Fig. S5), we employed the umbrella sampling method to calculate free energy profiles for the closure motion of the two RRMs. For the SUMO2/RBD/RNA triple complex, we used the same initial structure as for the previous RNA dissociation simulations. For the other combinations, we simply removed unneeded components from that structure to build initial structures. The structural similarity to the NMR structure of free RRMs, $Q_f$ was used as a biasing reaction coordinate. The biasing potential $V_{qbias}$ is mentioned above. For each system, we set 71 simulation windows with $Q_0$ evenly distributing from 0.1 to 0.8. Each simulation lasts $8 \times 10^6$ steps. The first $1 \times 10^6$ steps are treated as equilibration and are then discarded in later analysis. Another order parameter, PC0 is used for 2D-WHAM analysis[6]. 1D free energy profiles along PC0 axis are shown in Fig. S6.

### Efficiency of the shift of RNA-binding equilibration

1D Free energy profiles for RNA dissociation show that the difference of RNA-binding affinity of the SUMO2/RBD complex and that of the RBD by itself is around 2 kcal/mol. Therefore, an exchange equation for RNA-binding can be written down:

$$m \cdot CPEB + CPEB_S \rightarrow m \cdot CPEB_S + CPEB, \qquad \Delta G = -2kcal/mol$$

where $CPEB$ and $CPEB_S$ represent CPEB3 and SUMOylated CPEB3, and the prefix "$m\cdot$" represents RNA-bound complex. In equilibrium, we have:

$$\frac{[m \cdot CPEB_S]}{[m \cdot CPEB]} = e^{-\Delta G/RT} \frac{[CPEB_S]}{[CPEB]}$$

Based on Western blot experiments on CPEB3[7] and Orb2[8] extracted from the brain, the fraction of SUMOylated CPEB3, $[CPEB_S]_W$, is comparable to that of deSUMOylated CPEB3, $[CPEB]_W$, in the basal state. However, the Western blot result shows only the sum of $[m \cdot CPEB_S]$ and $[CPEB_S]$ or the sum of $[m \cdot CPEB]$ and $[CPEB]$, since the extraction and purification process would wash out most associated RNAs from RNA binding proteins[9]. Here we assume that CPEB3, as an efficient translational regulator, has much higher concentration than its target mRNAs. Given this assumption, $[m\cdot CPEB_S]$ or $[m\cdot CPEB]$ is negligible in comparison with $[CPEB_S]$ or $[CPEB]$, and thereby $[CPEB_S] \simeq [CPEB_S]_W$,

$[CPEB] \simeq [CPEB]_W$. Finally, we would find that the ratio of repressed target mRNA to active target mRNA, $\frac{[m \cdot CPEB_S]}{[m \cdot CPEB]}$, is approximately 30 in basal synapses.

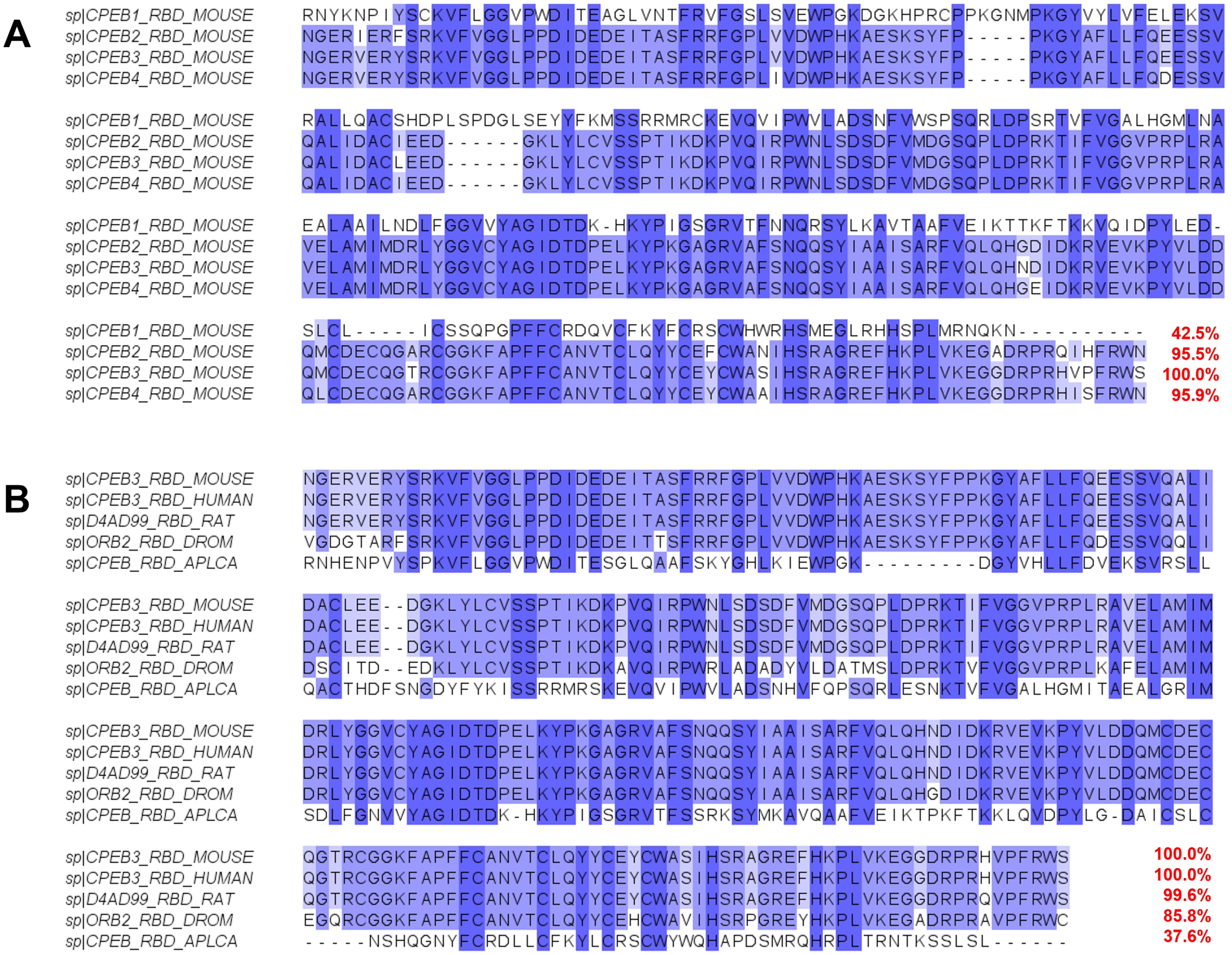


Figure 1: Multiple sequence alignment for the RNA-binding domain (RBD) of CPEB homologs. **A**. Four isoforms of mouse CPEB family. **B**. Homologs of mouse CPEB3, including CPEB3 in human, D4AD99 in rat, Orb2 in Drosophila and ApCPEB in Aplysia. For each residue, higher pairwise identity is marked by deeper blue color. Percentages of overall sequence identity are listed at the end of the alignment, comparing with mouse CPEB3.

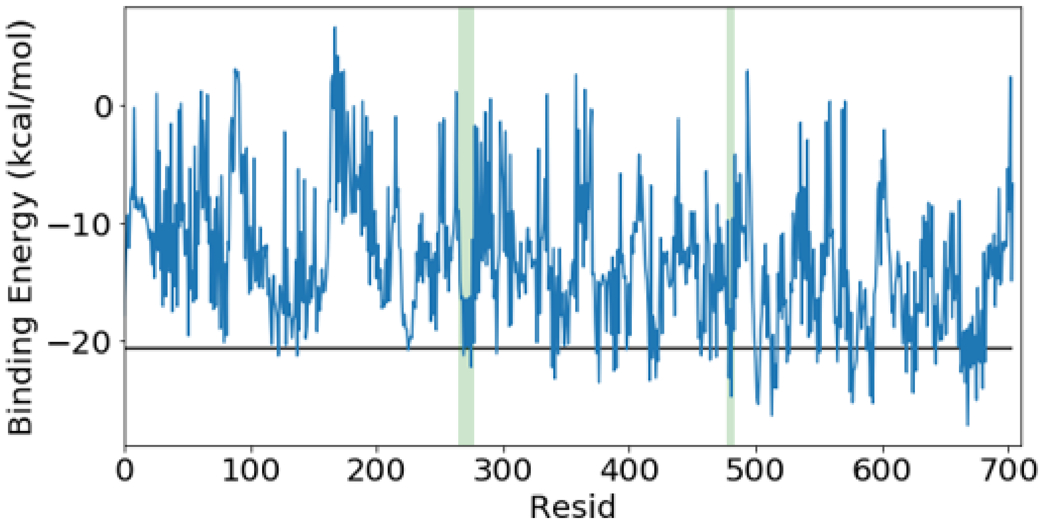


Figure 2: Binding energy of canonical SIM/SUMO2 complex structure when scanning the sequence of full length CPEB3 over the SIM peptide. The x axis is the residue index of the first amino acid in each CPEB3 peptide. The black line shows the binding energy for the original SIM peptide in the PDB structure (ID: 6JXW). The two green shaded regions highlighted peptides containing the predicted SIMs in CPEB3 via bioinformatic search (V273-V283; P484-W489).

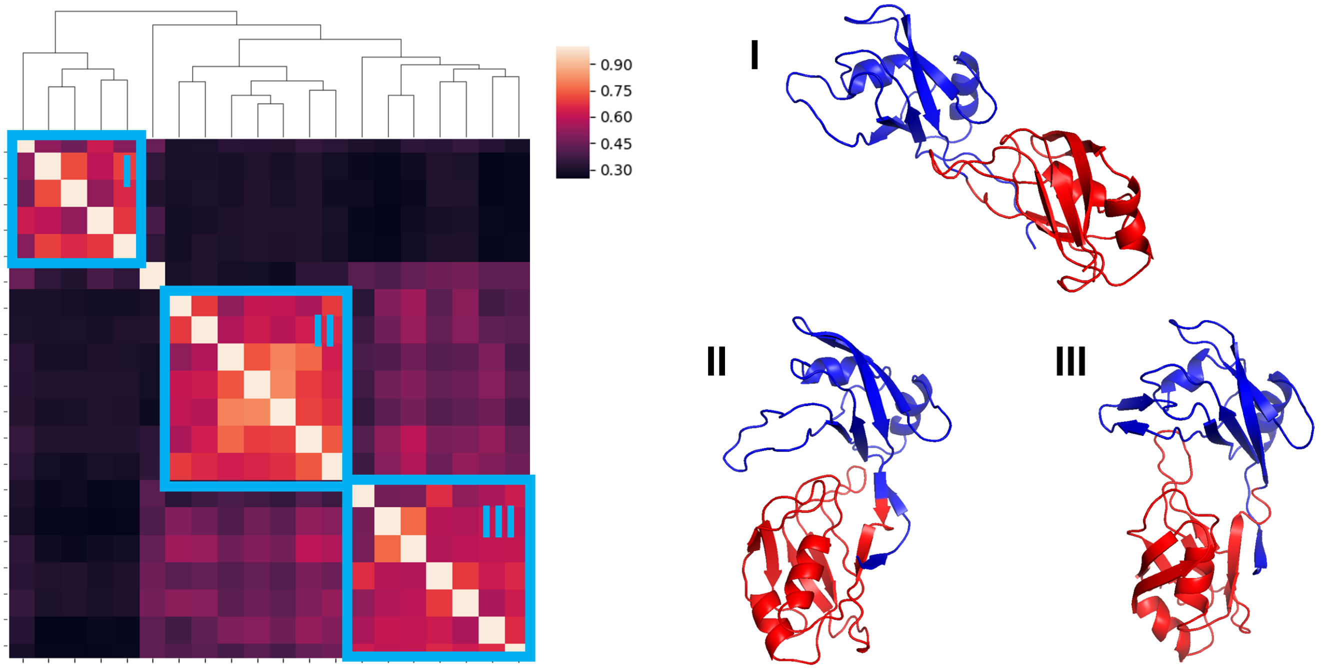


Figure 3: The 20 predicted structures of free RRMs can be divided into 3 clusters using mutual Q (only including inter-domain residue pairs) as the structural similarity metric. Representative structures for these 3 clusters are shown on the right.

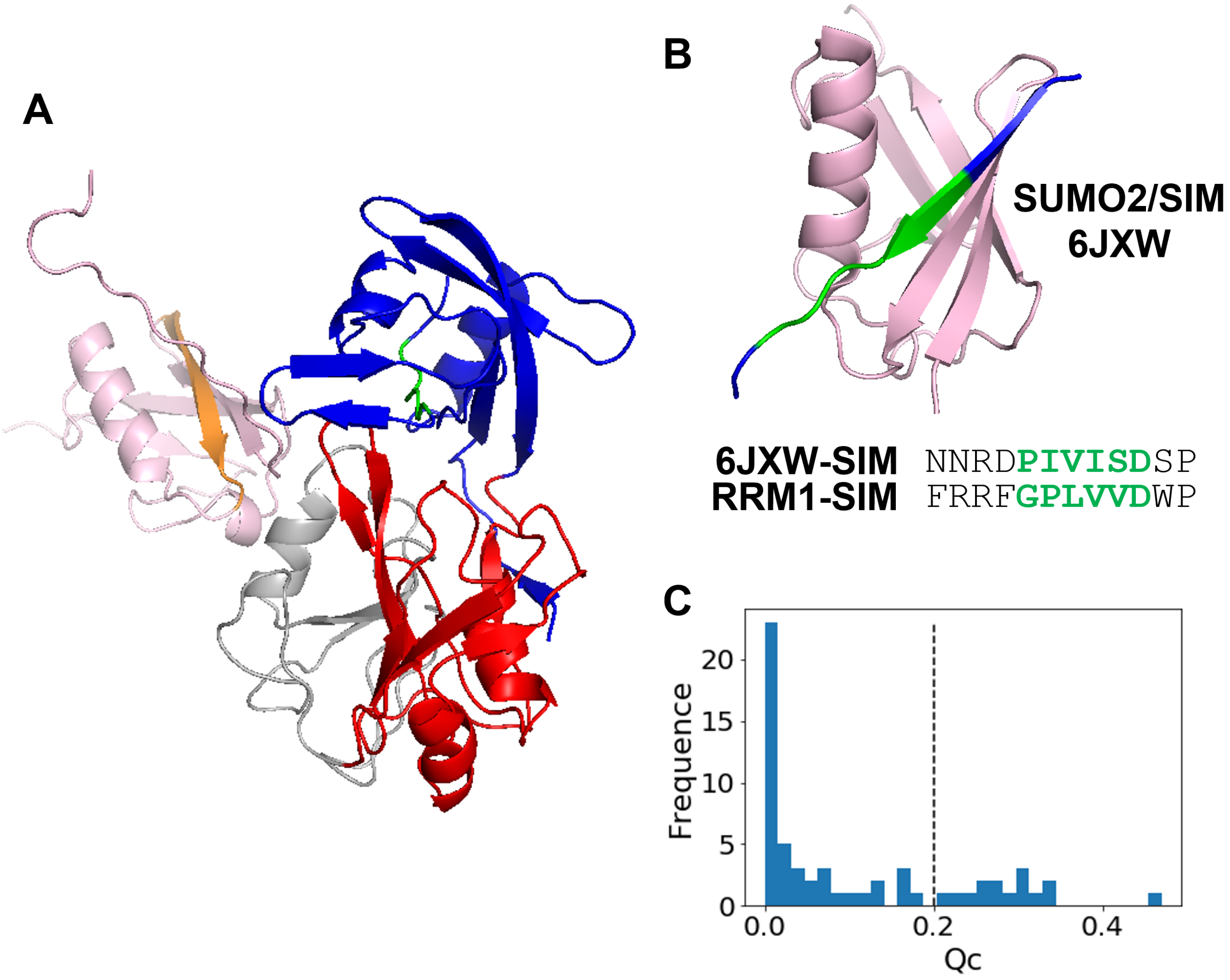


Figure 4: **A**. One example of the initial structure for the SUMO2/RBD complex structure prediction. The SUMO2 protein (pink) is docked into the ZnF domain based on PDB structure 2N1A. $\beta 2$ strand of SUMO2 is colored in orange and the RRM1-SIM is colored in green. **B**. Canonical SIM/SUMO2 complex (PDBID: 6JXW). SUMO2 is colored in pink. The SIM peptide is colored in blue, and the green fragment inbetween corresponds to the RRM1-SIM. The structure of SUMO2 and the green fragment is used as the reference to calculate the $Q_c$ value of SUMO2/RRM1-SIM in predicted SUMO2/RBD complex. **C**. The distribution of $Q_c$ value of SUMO2/RRM1-SIM in 60 predicted SUMO2/RBD complex structures. Structures with $Q_c$ larger than 0.2 is selected for further evaluation.

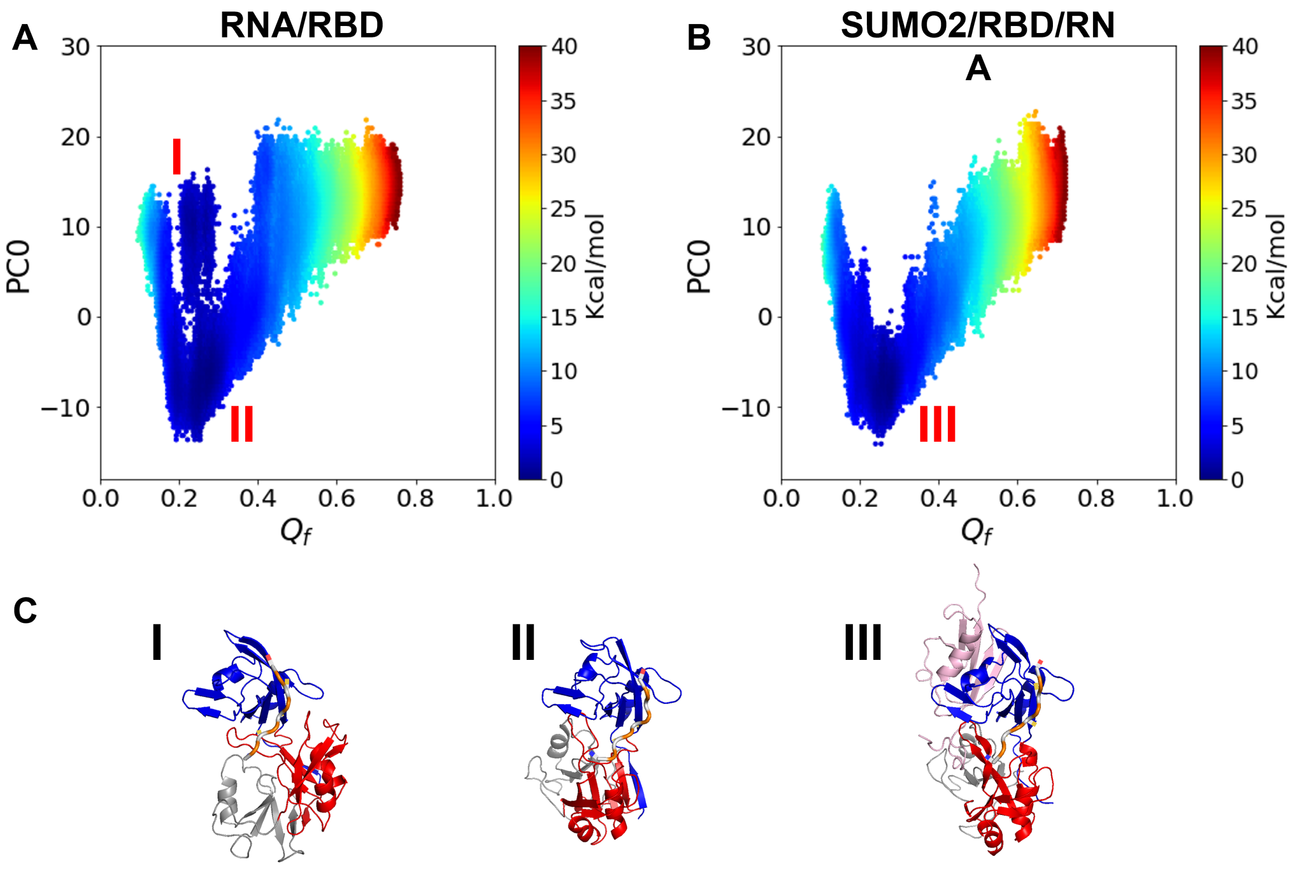


Figure 5: **A**. The free energy landscape for RBD/RNA using $Q_f$ and PC0 as the two order parameters. **B**. The free energy landscape for the SUMO2/RBD/RNA complex using $Q_f$ and PC0 as the two order parameters. **C**. Representative structures for the three free energy basins in Figure A and B.

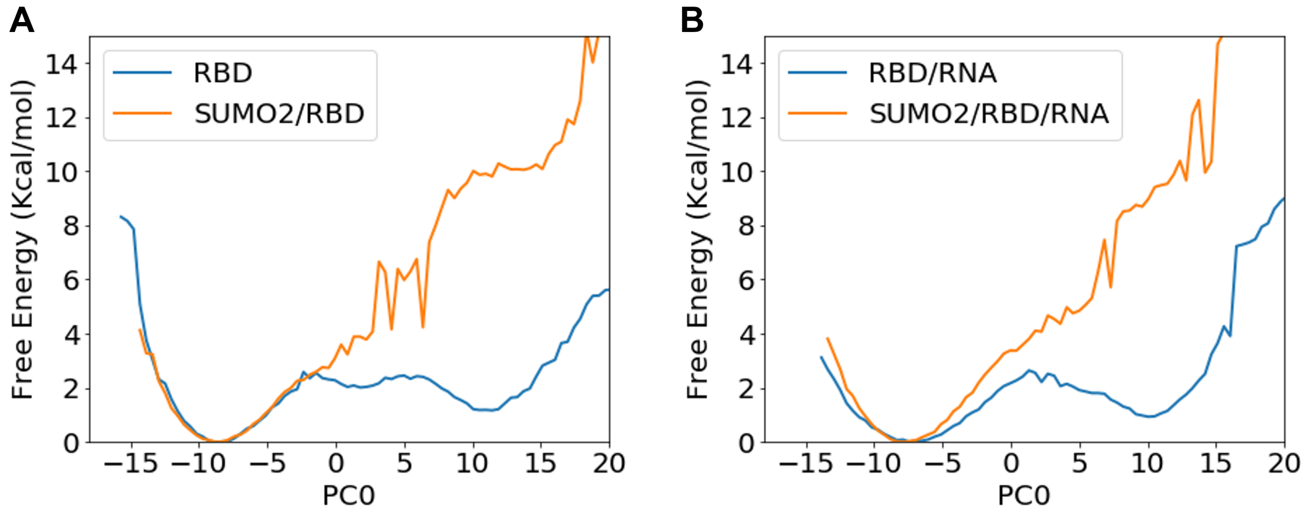


Figure 6: **A**. The 1D Free energy profile for RNA-free RBD and SUMO2/RBD complex using PC0 as the order parameter. **B**. The 1D Free energy profile for RNA-bound RBD and SUMO2/RBD complex using PC0 as the order parameter. The free energy of the open state (PC0 equals $10 \sim 30$) is significantly increased upon RBD binding with SUMO2 protein.

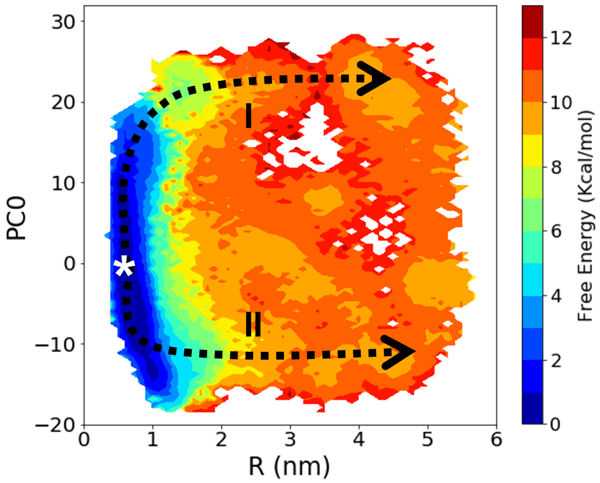


Figure 7: The two RNA dissociation pathways from RBD. These correspond with the pathway I and pathway II shown in Fig. 5B. The starting point of the dissociation pathways is marked by a white star.

### Supplementary Movies

**Movie S1. RNA dissociation pathway I from RRMs shown in Fig. 5B**

**Movie S2. RNA dissociation pathway II from SUMO2/RBD complex shown in Fig. 5B**

**Movie S3. RNA dissociation pathway I from RBD shown in Fig. S7**

**Movie S4. RNA dissociation pathway II from RBD shown in Fig. S7**

[6] Alan Grossfield. Wham: the weighted histogram analysis method, version 2.0.9.1. `http://membrane.urmc.rochester.edu/wordpress/?page_id=126`.

[7] Bettina Drisaldi, Luca Colnaghi, Luana Fioriti, Nishta Rao, Cory Myers, Anna M Snyder, Daniel J Metzger, Jenna Tarasoff, Edward Konstantinov, Paul E Fraser, et al. Sumoylation is an inhibitory constraint that regulates the prion-like aggregation and activity of cpeb3. *Cell reports*, 11(11):1694–1702, 2015.

[8] Ruben Hervas, Michael J Rau, Younshim Park, Wenjuan Zhang, Alexey G Murzin, James AJ Fitzpatrick, Sjors HW Scheres, and Kausik Si. Cryo-em structure of a neuronal functional amyloid implicated in memory persistence in drosophila. *Science*, 367(6483):1230–1234, 2020.

[9] Alexander G Baltz, Mathias Munschauer, Björn Schwanhäusser, Alexandra Vasile, Yasuhiro Murakawa, Markus Schueler, Noah Youngs, Duncan Penfold-Brown, Kevin Drew, Miha Milek, et al. The mrna-bound proteome and its global occupancy profile on protein-coding transcripts. *Molecular cell*, 46(5):674–690, 2012.
